# Self-organized pattern formation increases functional diversity

**DOI:** 10.1101/2020.09.27.315754

**Authors:** Janne Hülsemann, Toni Klauschies, Christian Guill

## Abstract

Self-organized formation of spatial patterns is known from a variety of different ecosystems, yet little is known how these patterns affect functional diversity of local and regional communities. Here we use a food chain model in which autotroph diversity is described by a continuous distribution of a trait that affects both growth rate and defense against a heterotroph. On a single patch, stabilizing selection always promotes the dominance of a single autotroph species. Two alternative community states, with either defended or undefended species, are possible. In a metacommunity context, dispersal can destabilize these states, and complex spatio-temporal patterns emerge. This creates varying selection pressures on the local autotroph communities, which feed back on the trait dynamics. Local functional diversity increases ten-fold compared to a situation without self-organized pattern formation, thereby maintaining the adaptive potential of communities in an environment threatened by fragmentation and global change.

## 1 Introduction

Biodiversity is a key prerequisite for the functioning of natural communities and ecosystems [Cadotte et al., 2011, Tilman, 1999, Hooper et al., 2005], the stable supply of ecosystem goods and services [Hooper et al., 2005] and the ability of communities and ecosystems to adapt to environmental change [Yachi and Loreau, 1999]. The maintenance of functional diversity within individual habitats may strongly rely on local processes and species interactions such as resource partitioning [Tilman, 1999] or neighborhood-dependent selection [Vasseur et al., 2011], which may generate sufficient niche differentiation among coexisting species. However, local communities are naturally embedded in a larger biogeographic context, and thus connected to other communities of adjacent habitats [Leibold et al., 2004]. The structure of such metacommunities may substantially influence the maintenance of functional diversity within local communities. A crucial aspect of this relationship is the connectivity among the different habitats, which is influenced by the permeability of the surrounding matrix and the dispersal abilities of the different species. For example, dispersal may have a positive influence on local functional diversity by allowing for source-sink dynamics (mass effects) where the establishment or persistence of local populations depends on the immigration of conspecific individuals from other localities [Leibold et al., 2004]. A necessary condition for this mechanism is spatial heterogeneity that allows differently adapted species to coexist on a regional scale [Amarasekare, 2003].

Most previous studies on the influence of dispersal on local population dynamics and functional diversity rely, however, on the assumption that spatial heterogeneity primarily results from variation in the abiotic environmental conditions. This premise neglects that spatial variation in species abundances may also emerge purely as a consequence of antagonistic (e.g. predator-prey) interactions of species in space that may lead to self-organized pattern formation (see Fig. 1a for an example) [Malchow, 1993]. The latter usually results from scale-dependent feedback, i.e., a situation where positive and negative feedback between species occurs at different spatial scales [Rietkerk and Van de Koppel, 2008]. For example, in many ecosystems, predators are more mobile than their prey [Peters, 1986], allowing them to suppress prey density over wider ranges (long-distance negative feedback), while prey only disperses locally. Positive feedback (self-facilitation and resource provisioning for the predator) therefore only acts over short distances and the prey might only be able to support a high predator density in confined areas.

**Figure 1:**
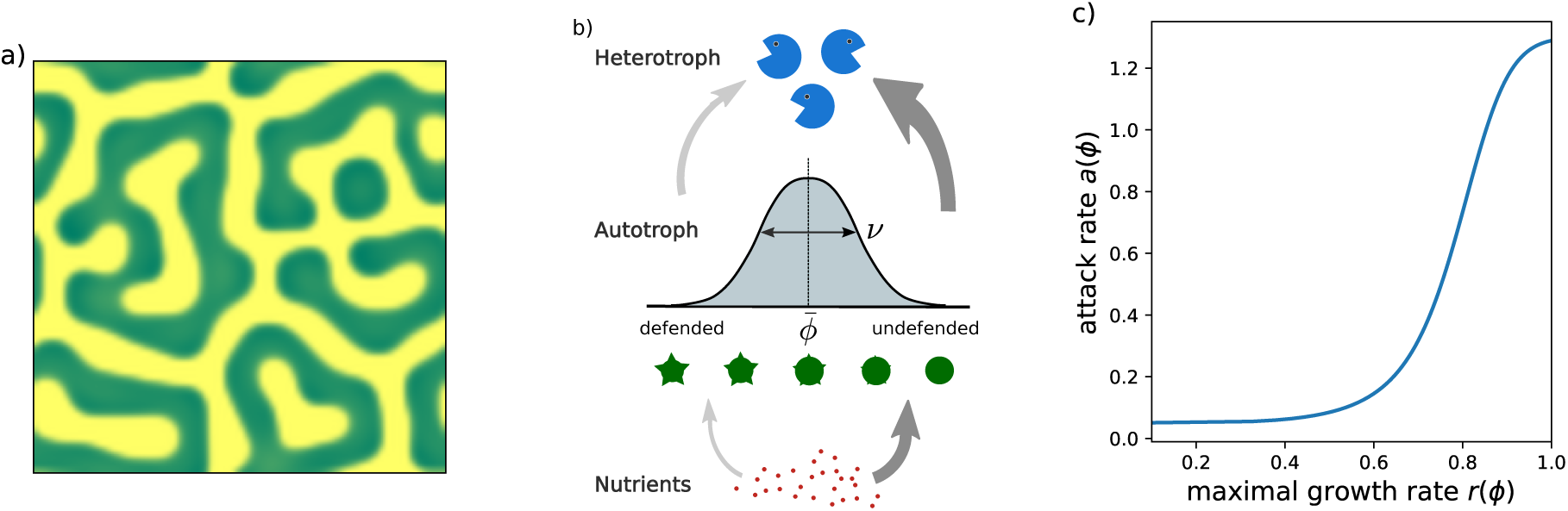
a) Example of a static Turing pattern. Here, vegetation (green) and bare soil (yellow) form a labyrinth-like pattern (created with the arid ecosystem model in [Rietkerk et al., 2002]). b) Conceptual representation of the local food chain model. The autotroph community exhibits a continuous (logit-normal) distribution of the trait *ϕ*, characterized by the mean 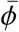 and the variance *ν*. A low trait value signifies a low attack rate of the heterotrophs (due to high investment into defense by the autotrophs), at the cost of a low maximal growth rate (visualised by thin arrows), whereas a high trait value leads to a high attack rate but also a high maximal growth rate (thick arrows). c) Shape of the trade-off between maximal growth rate *r*(*ϕ*) vs. attack rate *a*(*ϕ*), mediated by the trait *ϕ*.

Self-organized pattern formation through scale-dependent feedback was originally discovered and studied by Alan Turing [Turing, 1952]. According to his pioneering work, the development of spatial patterns originates from the destabilization of a spatially homogeneous state (i.e., where the abundances of the different species are the same in all habitats) by dispersal. Small, spatially non-uniform perturbations are thereby amplified and give rise to complex, but regular patterns in space. A necessary requirement for this so-called Turing instability is that dispersal rates are sufficiently high and differ among the species. The patterns can either be stationary like the spot- or stripe-patterns of some animal fur coats or, if generated by an oscillatory Turing instability, even vary in time (e.g. travelling waves). Traditionally, the mechanism proposed by Turing is only applied to spatially continuous systems, but the extension to networks of discrete habitat patches is well developed [Othmer and Scriven, 1971, Nakao and Mikhailov, 2010, Brechtel et al., 2018]. Spatial pattern formation due to a Turing instability has been found in a variety of ecological communities such as vegetation [Rietkerk and Van de Koppel, 2008], host-parasitoid [Hassell et al., 1994], plant-parasite [White and Gilligan, 1998], and plankton-fish systems [Medvinsky et al., 2002], and has been proposed as a potential explanation for patchy distribution of phytoplankton and herbivores in the oceans [Levin and Segel, 1976]. However, these previous studies only analyzed Turing instabilities in the context of pure biomass density patterns. Recently it as been shown that this can indirectly promote diversity by creating spatial niches for species with different environmental requirements [Cornacchia et al., 2018], but it remains open to what extent self-organized spatial pattern formation can directly influence the functional diversity of local communities and thus their adaptive potential to react to changes in the environment.

In general, we expect that locally, functional diversity is at least indirectly supported by spatial pattern formation. Spatially variable biomass densities of interacting species lead to heterogeneous biotic environments and thus to locally different selection pressures. Consequently, each habitat may possess a different species composition in the local community, which promotes regional species diversity. This, in turn, may feed back on the individual habitats and enhance the functional diversity of local communities, e.g via source-sink dynamics [Brown and Kodric-Brown, 1977, Shmida and Wilson, 1985]. Furthermore, we expect that the potential positive impact of spatial pattern formation on local functional diversity might be of particular relevance for systems where the prevailing selection regime is stabilising and would otherwise promote the dominance of a single species.

In this study, we investigate a food chain model with autotrophs and heterotrophs as prey and predators in a spatial context. We explicitly resolve the functional diversity of the autotrophs by considering a continuous trait distribution within each local habitat. A network of habitat patches, each hosting a local food chain, is interconnected via dispersal and thereby forms a metacommunity on the regional scale. Under the premise of suitable dispersal rates we expect to find spatial pattern formation in the biomasses, which may influence local functional diversity beyond classical source-sink dynamics.

## 2 Model description

The model consists of multiple food chains that are spatially distributed on interconnected habitat patches. The overall dynamics comprise local dynamics of biomasses and traits within each patch and spatial interactions (dispersal between patches).

### 2.1 Local biomass and trait dynamics

Within each habitat we consider the dynamics of a food chain consisting of nutrients (*N*), autotrophs (*A*) and heterotrophs (*H*) in a chemostat environment, with a diverse autotroph community. The diversity of the autotrophs is characterized by a logit-normally distributed trait *ϕ* (∈ [0 : 1]), which affects both defense level and maximal growth rate (Fig. 1b, c).

To describe the temporal changes in the shape of the underlying continuous trait distribution (Fig. 1b) we focus on its mean 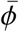 and variance *ν* by using the aggregate model approach [Norberg et al., 2001, Wirtz and Eckhardt, 1996]. Together with the nutrient concentration *N* and total biomass densities *A* and *H*, the full set of differential equations is

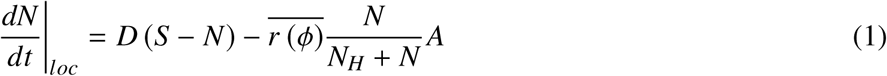

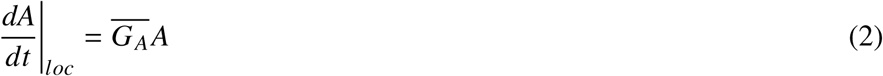

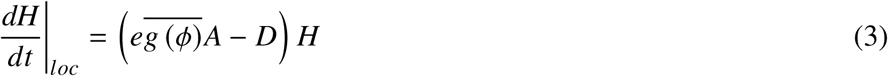

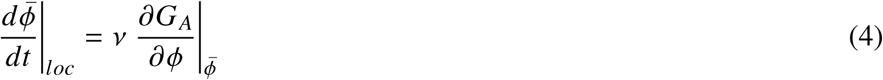

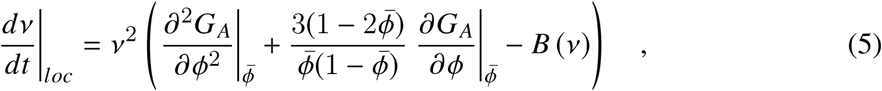

where |_*loc*_ means local dynamics without spatial interactions. The *per-capita* net growth rate (i.e. fitness function) of the autotrophs is

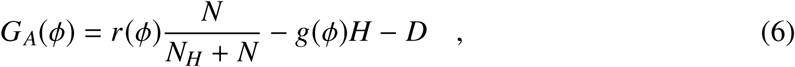

where the heterotroph’s grazing function is given by a Type II functional response

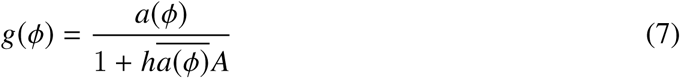

with a common handling time *h*. Following the aggregate model approach, the average of a function *f*(*ϕ*) (where *f* can be any of *r, G*_*A*_, g, or *a*) across the trait distribution is approximated up to the second order [Norberg et al., 2001, Klauschies et al., 2018]:

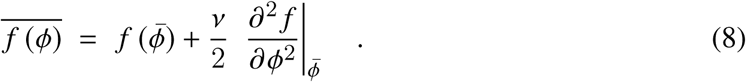

The nutrients *N* follow chemostat dynamics with supply concentration *S* and dilution rate *D*. The nutrient uptake by the autotrophs follows Michaelis-Menten kinetics, with half-saturation constant *N*_*H*_ and average maximal growth rate 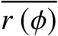. The autotrophs experience biomass losses due to grazing by the heterotrophs and the overall dilution rate *D*. The gain of heterotrophic biomass is determined by the average grazing rate 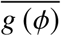 scaled by the conversion efficiency *e* and the total autotroph biomass *A*. The heterotrophs also experience losses due to dilution.

The direction and strength of natural selection that determines the change of the mean trait is given by the local fitness gradient 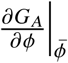. The speed of change is scaled by the trait variance *ν*, reflecting the functional diversity of the autotrophs: the presence of many different functional types (*ν* large) enhances adaptation capabilities of the trait, while a lack of functional diversity (*ν* small) slows down the adaptation process. Internal changes in *ν* are determined by the local shape of fitness landscape: its local curvature 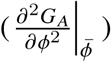 accounts for stabilizing or disruptive selection (reducing or enhancing *ν*, respectively), whereas the term including the local fitness gradient captures enhancing or reducing effects on *ν* through potential skewness of the trait distribution [Klauschies et al., 2018]. The latter keeps the mean trait in the range [0 : 1] by reducing *ν* (and thereby halting evolution in *ϕ*) when 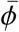 approaches the limits of the trait range. Potential effects of excess kurtosis and thus a bimodal shape of the trait distribution are neglected. For numerical reasons we included a boundary function *B*(*ν*) = *ϵ* /(*ν*_*max*_ − *ν*) to ensure that during transient dynamics the trait variance does not exceed the hypothetical maximal value of *ν*_*max*_ = 0.25 of a distribution on the interval [0 : 1]. The scaling parameter *ϵ* = 0.001 is sufficiently small to ensure that the boundary function has a negligible effect on the dynamics when the trait variance is not close to *ν*_*max*_.

### 2.2 Growth-edibility trade-off

The trait dynamics are driven by a trade-off between growth and defense of the autotroph [Tirok et al., 2011]. High values of *ϕ* denote high maximal growth rates *r* (*ϕ*) = (*r*_*max*_ − *r*_*min*_) *ϕ*^*s*^ + *r*_*min*_ (with shape parameter *s* < 1) but also high attack rates 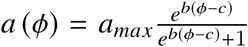, and vice versa (Fig. 1c). The parameter *b* determines the sensitivity of *a* to changes in *ϕ* and *c* sets the inflection point of the attack rate function. The trade-off between growth and edibility ensures that defended types (low *ϕ*) will thrive when predators are abundant, but are at a disadvantage when predator biomass is low.

### 2.3 Spatial interactions

In the spatial extension of our model each patch represents a local habitat *i*. For simplicity, we assume a linear arrangement of *n* patches with periodic boundary conditions, i.e., a ring. Nutrients and individuals move between neighbouring patches driven by differences in concentration or biomass density, respectively (diffusive movement). The local dynamics (Eqs. (1) to (5)) are thus supplemented by terms describing the effects of diffusion on nutrient concentrations, autotroph and heterotroph biomass densities (together summarized as *B*_*i*_), as well as on mean traits 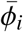 and trait variances *ν*_*i*_:

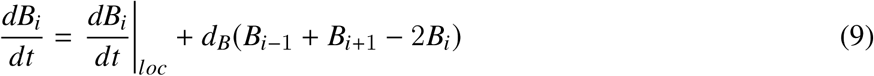

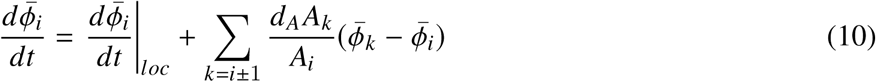

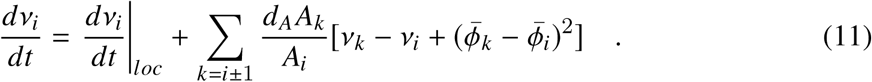

The parameters *d*_*B*_ (with *B* = *N, A*, or *H*) are the diffusion constants that describe the overall mobility of the respective component. If trait distributions differ between neighbouring patches, dispersal of the autotrophs also affects the evolution of mean traits 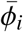 (Eq. (10) and trait variances *ν* (Eq. (11)) [cf. Norberg et al., 2001, Klauschies et al., 2018]. The evolution of the resident mean trait value is affected by the difference between mean trait values of resident and incoming biomass as well as by the ratio of incoming to resident biomass. Similarly, the evolution of the trait variance is affected by the difference between trait variances of resident and incoming biomass. Additionally, the variance increases if the mean trait of the incoming biomass differs from that of the resident, as this naturally broadens the resident trait distribution.

### 2.4 Turing instability

We here provide a very brief description of the procedure to determine whether a Turing instability occurs (and therefore the self-organized pattern formation can be expected). For more details of the mathematical derivation see [Othmer and Scriven, 1971, Brechtel et al., 2018] and the supplementary material, section S1. Whether or not the movement of nutrients, autotrophs and heterotrophs between local habitats lead to a Turing instability depends on the values of the diffusion constants *d*_*B*_. To determine whether a Turing instability occurs, one has to evaluate the eigenvalues *λ* of the *n* matrices

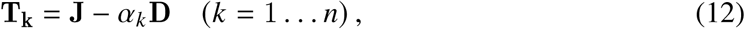

where **J** is the Jacobian matrix for the dynamics of *N, A*, and *H* evaluated at the fixed point of the local system, α_*k*_ are the eigenvalues of the Laplacian matrix (which describes the spatial network of the habitat patches) and **D** is a diagonal matrix composed of the diffusion constants *d*_*B*_. A Turing instability occurs if at least one of the *λ*’s of any of the **T**_**k**_ has a positive real part. If the dominant eigenvalues are a complex-conjugated pair, an oscillatory Turing instability (wave instability) occurs and spatio-temporal (i.e., non-stationary) patterns emerge. If the attractor of the local system is a limit cycle, one has to evaluate the Jacobian **J** along this limit cycle and calculate the Floquet multipliers *µ* of the **T**_**k**_. A Turing instability occurs if at least one of them has an absolute value > 1.

### 2.5 Simulation details and output variables

As key metrics influencing the pattern formation process we varied the diffusion constants *d*_*N*_, *d*_*A*_, and *d*_*H*_. The other parameters were set to constant values oriented at planktonic systems (Tab. 1). For each combination of diffusion constants, 100 replicates with random initial conditions were calculated 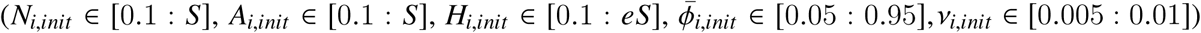. As measures for the local and regional functional diversity we calculated the mean of *ν* and the variance of 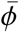 across patches and replicates, based on the temporal averages of the respective variables per patch. Time series were first simulated over 100 000 time steps to eliminate transient dynamics and then for another 15 000 time steps to evaluate temporal means of *ν* and 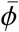. Additionally, we calculated the probability of maintaining local or regional functional diversity as the fraction of replicates with mean (*ν*) > 10^−5^ and 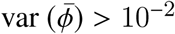. Values below these thresholds usually tended towards zero, but at a very slow rate.

The solutions of the differential equations were obtained numerically in C using the BDF method of the SUNDIALS CVODE solver [Hindmarsh et al., 2005] with relative and absolute tolerances of 10^−15^. The output data were studied using Python 3.7.3 and several Python packages, in particular NumPy version 1.16.2 and Matplotlib version 3.0.3 [Hunter, 2007, Van Der Walt et al., 2011].

## 3 Results

### 3.1 Local dynamics

We first describe the dynamics of the local food chain with trait dynamics (Eqs. (1)-(5), Figs. S2 and S3 in the supplementary material). An important feature of this model is that it exhibits bistability: one of the attractors is characterized by low trait values 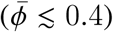 and thus a high defense level of the autotrophs (henceforth referred to as the defended attractor), while the other is characterized by very high trait values of 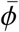close to 1, entailing a low defense level and a high growth rate of the autotrophs (from now on referred to as the undefended attractor). The bistability emerges due to the shape of the trade-off between growth rate and edibility (Fig. 1c): On both attractors, changing the trait in either direction increases costs (increasing attack rate or decreasing growth rate) more than benefits (increasing growth rate or decreasing attack rate). On the defended attractor, the autotroph biomass is high because the strong defense leads to a low heterotroph biomass. The autotrophs are bottom-up controlled by a low nutrient concentration. Conversely, on the undefended attractor, the autotrophs are under top-down control and their biomass is kept at a lower level than on the defended attractor, while nutrient concentration and heterotroph biomass reach higher levels.

**Table 1:** Description and values of the model parameters.

| parameter | description | dimension | value |
| --- | --- | --- | --- |
| $S$ | substrate supply concentration | $\text{mass} \cdot \text{area}^{-1}$ | 4.8 |
| $D$ | dilution rate | $\text{time}^{-1}$ | 0.3 |
| $N_H$ | half-saturation constant for nutrient uptake | $\text{mass} \cdot \text{area}^{-1}$ | 1.5 |
| $r_{max}$ | maximum growth rate of the autotroph | $\text{time}^{-1}$ | 1 |
| $r_{min}$ | minimum growth rate of the autotroph | $\text{time}^{-1}$ | 0.1 |
| $s$ | shape parameter of the growth rate function | - | 0.3 |
| $a_{max}$ | maximum attack rate of the heterotroph | $\text{area} \cdot (\text{mass} \cdot \text{time})^{-1}$ | 1.3 or 2.0 |
| $b$ | scaling parameter of the attack rate function | - | 8 |
| $c$ | scaling parameter of the attack rate function | - | 0.4 |
| $h$ | handling time for the autotrophs | time | 0.53 |
| $e$ | conversion efficiency of the heterotroph | - | 0.33 |
| $\epsilon$ | scaling parameter of the boundary function | - | $10^{-3}$ |
| $v_{max}$ | maximum value of the trait variance | - | 0.25 |

The nature of the undefended attractor depends on the maximum attack rate *a*_*max*_. At low values of *a*_*max*_, the attractor is a stable fixed point, but at *a*_*max*_ ≈ 1.35 it undergoes a Hopf bifurcation and a limit cycle with oscillations in the nutrient concentration and the biomass densities occurs. The defended attractor, conversely, is always a stable fixed point that is only marginally affected by the variation of *a*_*max*_. On both attractors and irrespective of the Hopf bifurcation, the mean trait 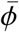 settles at a constant value. Initially, selection can be disruptive, but eventually the trait variance *ν* always slowly decays to 0 due to stabilizing selection closer to the respective attractor, thereby preventing any further temporal change of 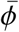.

### 3.2 Spatial dynamics

For the analysis of the spatially extended system we first established ranges of the diffusion constants that lead to a Turing instability (TI). These ranges differ for the two attractors, with the diffusion constants leading to a TI of the defended attractor being a subset of those that lead to TI of the undefended attractor (supplementary material, Fig. S1). We also established that the leading eigenvalues of the matrices **T**_**k**_, Eq. (12), are always complex, i.e., an oscillatory TI occurs. There are thus three different situations: no TI of any attractor, TI of only the undefended attractor, and TI of both attractors. However, as diffusion constants for the third case have to be very high, the strong coupling between the patches homogenizes the system: spatio-temporal patterns in the biomass densities still occur, but both local and regional functional diversity are almost always lost as all patches settle on a single attractor. We therefore do not consider this case. The occurrence of a Hopf bifurcation and a Turing instability of the undefended attractor thus leaves us with four qualitatively different cases: 1) undefended attractor is a fixed point and no TI occurs, 2) undefended attractor is a limit cycle and no TI occurs, 3) undefended attractor is a fixed point and a TI occurs, and 4) undefended attractor is a limit cycle and a TI occurs.

Fig. 2 shows representative time series for the four cases in a system with *n* = 6 patches. Local functional diversity (trait variance *ν*) was maintained in all four cases (Fig. 2i to l), even without TI (cases 1 and 2). In these first two cases, trait variance does not vanish because coupling of patches with defended and undefended autotroph communities, respectively, ensures that the potential loss of maladapted phenotypes within a single patch is counterbalanced by their continuous immigration from a different patch (last term in Eq. (11)). In case 2, weak diffusive coupling also tends to synchronise the biomass oscillations on patches that host an undefended autotroph community (Fig. 2b). However, mean trait values 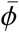 remain constant, as the low value of trait variance *ν* does not allow for a tracking of the varying selection pressure due to the comparably fast biomass oscillations.

**Figure 2:**
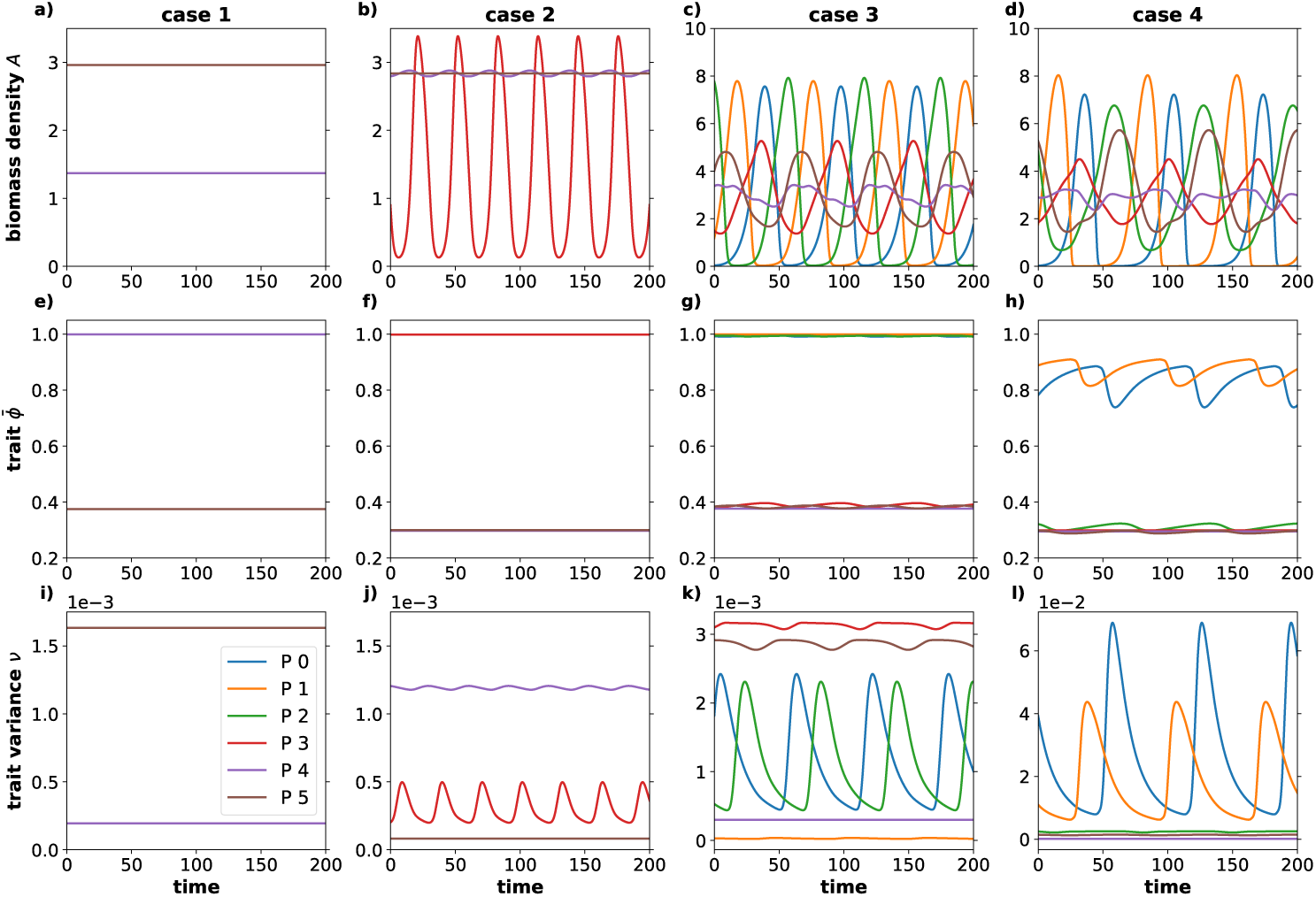
Exemplary time series of autotroph biomass densities (top row), mean traits 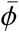 (middle row) and trait variances *ν* (bottom row, y-axes are scaled with the number at the top left corner of the panels) for spatial systems in cases 1 to 4 with 6 patches. In cases 1 and 2, *d*_*N*_ = 10^−2^, in cases 3 and 4, *d*_*N*_ = 1. In all four cases, *d*_*A*_ = 10^−5^ and *d*_*H*_ = 10^−3^.

In cases 3 and 4 (TI of the undefended attractor) complex spatio-temporal dynamics are seen (Fig. 2c and d). On a given patch, the biomass oscillations are notably slower than those induced by the Hopf bifurcation, their amplitudes are larger, and they now also affect the patches with a defended autotroph community. The maintained level of trait variance *ν* is markedly higher than in the cases without TI, which allows for a faster tracking of varying selection pressures and leads to visible temporal dynamics in the mean trait 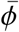. These results rely on the presence of both the defended and the undefended attractor in the metacommunity. If only one of the attractors is present, pattern formation in the nutrient concentration and biomass densities still occurs, but both local and regional functional diversity are lost over time (supplementary material, Fig. S4).

The diffusion constant of the autotrophs (*d*_*A*_) directly affects the trait dynamics within the metacommunity (cf. Eqs. 10 and 11). Fig. 3 shows the effect of *d*_*A*_ on the local (Fig. 3a) and regional diversity (Fig. 3b) as well as on the probability of preserving them (Fig. 3c and d). Increasing *d*_*A*_ positively affects local diversity, but has a neutral to negative effect on regional diversity. The latter effect is due to mean traits of communities on different patches becoming more similar when spatial coupling between the patches intensifies. Regarding local diversity, we observed clear and consistent differences between the four cases. Without TI (cases 1 and 2), local diversity was lowest and similar for the two cases. Case 3 exhibits an elevated level of local diversity, especially towards higher values of *d*_*A*_, and case 4 stands out by showing an increase in local diversity by about an order of magnitude compared to the cases without TI. However, case 4 also exhibits the largest loss of regional diversity (mean traits of the communities on different patches become more similar as *d*_*A*_ increases) and the probability of retaining local and regional diversity in the metacommunity strongly declines when *d*_*A*_ is larger than 10^−5^. (Note that mean local and regional diversity, Fig. 3a and b, was only calculated using the replicates in which the respective diversity measure was retained.)

**Figure 3:**
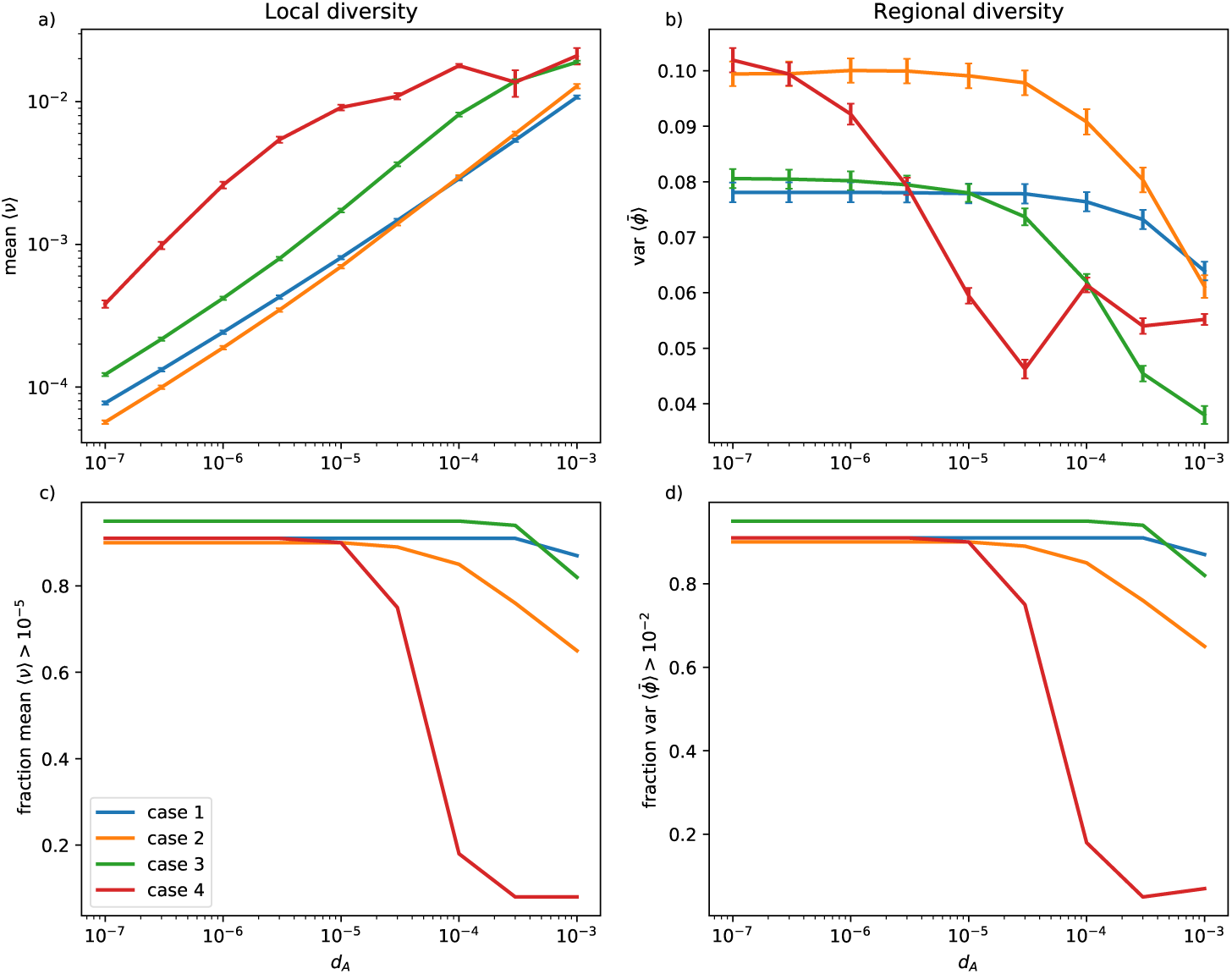
The degree of trait diversity maintenance depends on the mobility of the autotrophs. In panel a) the mean trait variance *ν* across 6 patches is shown as a measure of local diversity, in b) the variance of mean trait values 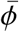signifies regional diversity. Results were obtained as arithmetic means over 100 simulation runs with randomized initial conditions, error bars denote standard errors. In panels c) and d), the fraction of simulation runs in which local (c, mean(*ν*) > 10^−5^) or regional 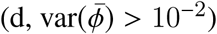 diversity were maintained, are shown.

Last, we demonstrate that the spatio-temporal pattern formation induced by the Turing instability is indeed the driving force behind the increased level of local diversity. Comparing the maximal eigenvalue or Floquet multiplier (depending on whether the undefended attractor of the local dynamics is a fixed point or a limit cycle) of the matrices **T**_**k**_, which determines whether a TI occurred, with the level of preserved local diversity (trait variance *ν*) over a wide range of values of the diffusion constants *d*_*N*_ and *d*_*H*_ shows that both metrics are coherently increasing with the distance of the diffusion constants from the Turing instability (Fig. 4). This visual inspection of their dependency is also quantitatively supported (Spearman rank-correlation *r*_*s*_ = 0.86 for stable local dynamics, and *r*_*s*_ = 0.84 for oscillatory local dynamics. See supplementary material, section S5, for additional information).

**Figure 4:**
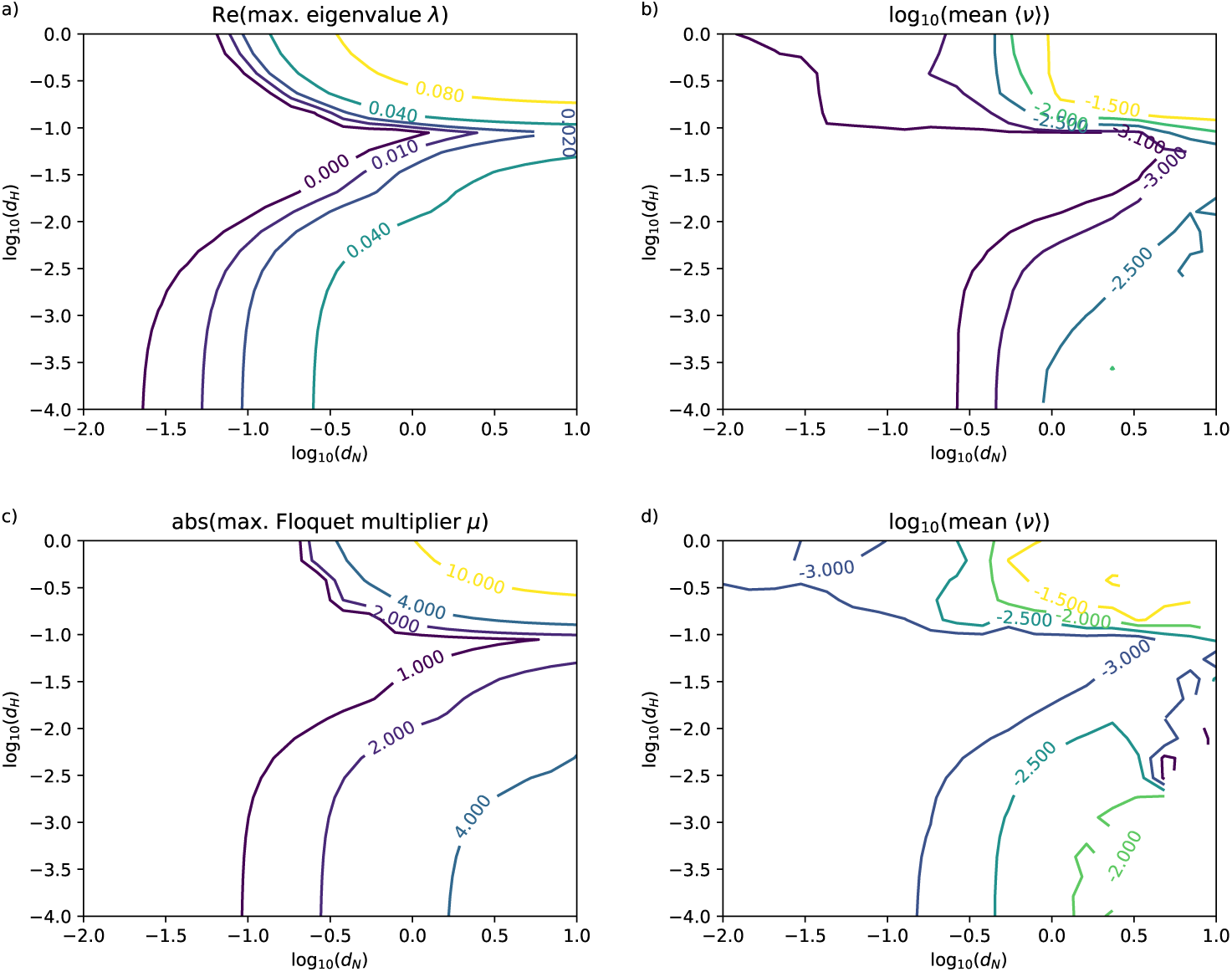
Correspondence between Turing instability and local diversity. Panels a) and c) show the maximum eigenvalue and Floquet multiplier, respectively, of the matrices **T**_**k**_ (Eq. (12)), panels b) and d) show on a logarithmic scale the mean trait variance (averaged over patches and simulation runs), .i.e., the local diversity. For a) and b), the undefended attractor is a fixed point (*a*_*max*_ = 1.3), for c) and d) it is a limit cycle (*a*_*max*_ = 2.0). In both cases, *d*_*A*_ = 10^−5^.

## 4 Discussion

Preserving functional diversity is essential for the functioning of ecosystems [Tilman, 2001, Hooper et al., 2005] and their persistence in the face of accelerated global change [Duraiappah et al., 2005]. Recent studies have shown that locally (i.e., in a specific habitat), its maintenance may strongly rely on the immigration of novel phenotypes [Norberg et al., 2001, Norberg, 2004, Klauschies et al., 2018]. However, these studies do not explicitly account for the spatial context and, thus, potential interactions between regional and local functional diversity. Here, regional functional diversity refers to the existence of different mean values of the distribution of a relevant functional trait throughout a metacommunity. In the present study, we overcome this limitation by investigating the potential impact of dispersal between different habitats on the functional diversity of local communities through a novel mechanism based on self-organized pattern formation. For a wide range of dispersal rates, spatio-temporal patterns in the biomass densities may form, which feed back on the distribution of the functional trait. If these local biomass-trait feedbacks interact with regional diversity, local functional diversity increases up to tenfold compared to scenarios without self-organized pattern formation.

### 4.1 No or low dispersal between patches

If no dispersal between habitat patches occurs, stabilising selection leads to the loss of functional diversity in the isolated local communities. Either fast-growing, but completely undefended autotrophs, or slow-growing, well defended types, dominate in the individual communities, but coexistence of the two strategies in one patch is not possible. Over time, these isolated communities thus lose their ability to adapt to changing environmental conditions, which makes them susceptible to perturbations [Gunderson, 2000]. Regional functional diversity, on the other hand, is not directly affected by the absence of spatial interactions.

At low dispersal rates that do not lead to self-organized pattern formation, a minor reduction in regional functional diversity occurs as dispersal of individuals with different trait values through the landscape causes the mean traits of defended and undefended communities to converge slightly. At the same time, however, the continuous exchange of individuals with different trait values between the different habitats also slightly enhances local trait diversity. This is consistent with source-sink dynamics, where the maintenance of local diversity results from dispersal of species from favourable to unfavourable habitats [Shmida and Ellner, 1984, Pulliam, 1988]. Species that cannot maintain a permanent, self-sufficient population locally can still establish a small population with the help of recurring immigration [Mouquet and Loreau, 2002, Shmida and Wilson, 1985]. This increases the local range of functional types (i.e., local functional diversity) in comparison to isolated local habitats. Dispersal thus enhances local diversity but retains sufficient differences between local habitats to ensure regional diversity. However, immigrating species from habitats with differing conditions will always stay maladapted and the persistence of their population in the target patch remains dependent on ongoing immigration.

### 4.2 Self-organized pattern formation at intermediate dispersal rates

When the dispersal rates of the predators and prey as well as the diffusion rate of the nutrients are sufficiently high to induce a Turing instability, spatio-temporal patterns in the nutrient concentration and biomass densities form. These patterns create differences in per capita predation rates and nutrient availability, and thus different selection pressures on the autotroph communities that are fluctuating both in space and time. However, with no independent source of regional diversity, e.g. due to the coexistence of defended and undefended communities in the landscape, these fluctuations in selection pressures proved not strong enough or to vary too quickly to overcome the general stabilising selection regime that over time eliminates local diversity.

However, if there is an additional source of regional diversity that is independent of the self-organized pattern, local diversity is maintained on a level that is far beyond what could be expected from simple source-sink dynamics. The fluctuating selection pressure together with the influx of external phenotypes is now sufficient to maintain local diversity on a level that allows for continuing adaptation of the local communities. We suggest the following explanation: When the mean trait of a local community is at its optimal value (maximising the fitness function *G*_*A*_), the curvature of the fitness landscape is necessarily negative and trait variance (local diversity) tends to decrease. Under a fluctuating selection regime, however, the optimal trait value varies over time, which allows different phenotypes to thrive at different moments in time. Furthermore, the immigrating species will usually be maladapted, which in combination with the constantly changing biotic environment keeps the mean trait of the local community sufficiently far from its current optimal value that the curvature of the fitness landscape may be less negative or even positive). Thereby, local dynamics do not reduce trait variance as strongly and can be counter-balanced by the direct increase of trait variance by the immigration of species with different trait values. When additional temporal variability in the form of a limit cycle of the local population dynamics is present, local diversity is increased even further. This demonstrates that local diversity can arise from a complex interaction between processes on local and regional scales.

We hypothesize that even without an independent source of regional diversity it is possible that self-organized pattern formation and the entailing varying selection pressures induce permanent local diversity. First evidence for this is provided by Eigentler and Sherratt [2020], who analyze the coexistence of two plant species in a savanna vegetation model. In general, conditions under which pattern formation might facilitate local diversity include a less strongly stabilising selection regime, which would enable higher degrees of freedom in the evolution of the autotrophs, a more flexible shape of the local trait distributions (e.g., assuming a beta distribution instead of a transformed normal distribution [Klauschies et al., 2018]), or stationary instead of oscillatory Turing patterns (such as those in Fig. 1a). These stationary patterns would create selection pressures that only vary between habitats, but are constant over time in each habitat. This would allow the local communities to slowly adapt to the prevailing conditions and thereby create a certain amount of regional diversity in a self-organized manner, which in turn could increase local functional diversity via source-sink dynamics.

### 4.3 High dispersal rates

The amount of local functional diversity maintained correlates strongly with the strength of the Turing instability, i.e., the more pronounced the spatio-temporal patterns are, the higher is the local functional diversity (Fig. 4). Generally, this happens when the diffusion or dispersal rates increase, which leads to a higher exchange of nutrients, biomass, and especially different phenotypes between the habitat patches.

At the same time, however, increasing dispersal rates can also reduce functional diversity by causing a progressive homogenization of the metacommunity. In fact, stronger ecological coupling may lead to a decrease in regional diversity [Mouquet and Loreau, 2003], which in turn lowers local diversity: phenotypes that are exchanged between local habitats are more and more similar, which reduces the stimulating effect of dispersal on local diversity. In our system, this is of particular relevance since regional variability relies on the coexistence of defended and undefended communities in the landscape. Progressive homogenization thus has a strong effect since it increases the probability for all local communities to adopt the same strategy (fast growing, undefended, or slow growing, well defended), corresponding to a complete loss of regional diversity.

### 4.4 Applicability in different ecosystem types

While we developed our model with a planktonic system in mind, the wide range of autotroph dispersal rates that allows for self-organized pattern formation (Fig. 3) suggests that our findings may apply to a variety of different ecosystems. These range from aquatic environments with relatively mobile autotrophs (mid to upper end of the *d*_*A*_ spectrum) to terrestrial ecosystems with often very limited autotroph dispersal (lower end of the *d*_*A*_ spectrum).

In aquatic metacommunities, local habitats are given by ponds or lakes that are interconnected by dispersal, e.g. through overflows and rivulets [Leibold et al., 2004, Vanormelingen et al., 2008]. Furthermore, aquatic metacommunities can establish in larger waterbodies like large lakes, seas and oceans [Leibold and Norberg, 2004] where local communities form in confined areas like embayments or gyres and are interconnected by water currents. The autotroph level mainly consists of phytoplankton which behaves like particles in water and exhibits moderate mobility. Inorganic nutrients are considerably smaller particles with diffusion rates several orders of magnitude higher than the diffusion rates of the phytoplankton. Due to active movement herbivorous zooplankton can also achieve much higher dispersal rates than the phytoplankton. Previous theoretical studies showed that self-organized pattern formation can occur in plankton systems [Malchow, 1993, Medvinsky et al., 2001, 2002, Ruan, 1998]. However, to our knowledge, none of these studies considered pattern formation as an explanation for plankton diversity and as a potential mechanism contributing to resolving the paradox of the plankton [Hutchinson, 1961, Scheffer et al., 2003].

This mechanism of course will always interact with other mechanisms creating diversity in aquatic communities. Depending on the scale considered, even small systems like ponds can be internally structured with clear distinctions between littoral, pelagic, and benthic zones. Furthermore, vertical stratification in lakes creates a structuring with sufficiently different environmental conditions that allow different communities to develop. On the other hand, seasonal mixing events temporally equalise diffusion rates, thereby eliminating a necessary condition for self-organized pattern formation.

In terrestrial systems the autotroph level consists of sessile plants, which move only once in their lifetime during seed or propagule dispersal, while nutrient transportation (e.g. via ground water) and active herbivore movement are once again relatively fast. The dispersal rates of plants are consequently determined by their generation time as well as their dispersal strategy [Pueyo et al., 2008]. Since both factors can vary greatly between terrestrial plant species, a variety of different autotroph dispersal rates exist in terrestrial metacommunities, which are covered by the results in this study. Species diversity in terrestrial ecosystems is explained by a number of classical theories like the intermediate disturbance hypothesis [Connell, 1978] or the Janzen-Connell-hypothesis [Hyatt et al., 2003]. The former links plant diversity to non-equilibrium dynamics, while the latter traces it back to an interaction between a clumped seed dispersal around parent trees and seed survival depending on propagule density and distance from the parent tree. Self-organized pattern formation, which was found in several terrestrial environments like arid ecosystems [Kéfi et al., 2008, 2010], savanna grasslands and “ribbon forests” (striped patterns of tree lines) [Rietkerk and Van de Koppel, 2008], might be a valid addition to these well-established theories.

### 4.5 Summary and conclusions

In line with previous studies [Venail et al., 2008, Mouquet and Loreau, 2003], our findings show that dispersal acts as a link between local and regional diversity and that it can have both positive and negative effects on species coexistence in a metacommunity. Besides having the potential to cause self-organized pattern formation, dispersal determines to what extent functionally different species are exchanged between local habitats and therefore how regional diversity affects local diversity. Low dispersal rates that do not initiate pattern formation enhance local diversity only marginally via source-sink dynamics. At the other end of the spectrum, strong dispersal tends to homogenize the landscape and thereby eliminates any differences between the habitats, which ultimately also leads to the loss of functional diversity in local communities. In contrast, intermediate dispersal leads to the highest amount of preserved diversity. It enhances local functional diversity through an interplay of source-sink dynamics with variable selection pressures caused by self-organized pattern formation while preserving regional differences in the local trait distributions.

This study is the first to explicitly show how self-organized pattern formation by the Turing mechanism increases functional diversity in a general ecosystem model, thereby providing an additional explanation for local diversity in various ecosystems. To fully understand spatial and temporal variation of biodiversity we thus argue that it is necessary to take regional processes like pattern formation and their effects on local dynamics into account. Since self-organized pattern formation occurs only for certain combinations of dispersal rates, factors like obstacles or corridors that reduce or enhance the flux of nutrients and organisms in nature potentially have a strong effect on the diversity of natural metacommunities. This has important implications for decision making in ecological management, as both facilitating or hindering dispersal can have adverse effects on pattern formation and thus the maintenance of biodiversity.

## Acknowledgements

This study was partly supported by the German Research Foundation (DFG) under grant number GU 1645/1-1. The authors thankfully acknowledge helpful comments by Ellen van Velzen and Ursula Gaedke on an earlier version of the manuscript.

## Conflict of interest

The authors declare no conflict of interest.

## Author contributions

CG and TK conceived the study design. JH and CG wrote the computer code, performed the numerical simulations and evaluated the data. All authors interpreted and discussed the results. JH wrote the first draft of the manuscript (with contributions of CG), CG led the editing. All authors contributed critically to the drafts and gave final approval for publication.

## Supplement

### S1: Turing instability in spatially discrete systems

In this section we briefly outline the central ideas and the most important mathematical steps of the Turing mechanism that leads to self-organized pattern formation. We first describe the more commonly considered case of continuous space (much more detailed descriptions of which can be found in many advanced textbooks of theoretical biology, e.g. [Murray, 2003]) and then make the step to spatially discrete systems. For simplicity, and for consistency with the main text, we only consider one spatial dimension with coordinate *x*. The extension to more complex scenarios is straight forward and does not change the core of the mathematical considerations.

In our model we assume that nutrient molecules as well as herbivore and plant individuals move randomly in space and that net fluxes are only driven by differences in their concentration or biomass densities, respectively (as opposed to e.g. active searching of predators for regions with high prey density). In a spatially continuous setting, such movement is conceptualized as random diffusion and is mathematically described by Fick’s law that relates the temporal change of a concentration or biomass density to its second derivative in space, scaled by a diffusion constant. (In this appendix we therefore refer to any random movement to diffusion, even though the active movement of the herbivores is more accurately described as random dispersal.)

Since nutrients, plants, and herbivores also interact with each other at every point in space (uptake of nutrients by the plants, consumption of plants by the herbivores), they form a so-called reaction-diffusion system:

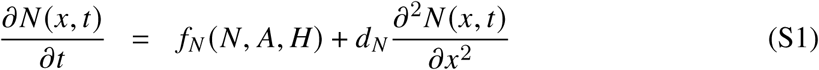

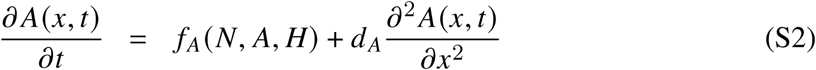

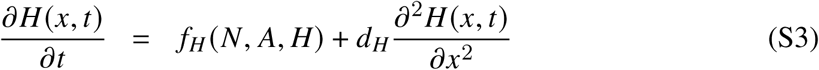

where *f*_*i*_ (*i* = *N,A,H*) is the total rate of change due to local interactions and *d*_*i*_ is the diffusion constant of each component. Note that we do not have to consider the dynamics of mean trait and trait variance here. While they are certainly affected by diffusion, they do not represent independent physical objects that can move through space.

In his seminal paper, Turing [1952] formulated the idea that such a system can exhibit a so-called diffusion-driven instability (or Turing instability): If the system possess a homogeneous steady state (*N*_*h*_,*A*_*h*_,*H*_*h*_), for which *f*_*N*_(*N*_*h*_,*A*_*h*_,*H*_*h*_) = *f*_*A*_(*N*_*h*_,*A*_*h*_,*H*_*h*_) = *f*_*H*_(*N*_*h*_,*A*_*h*_,*H*_*h*_) = 0 holds, that is stable in the absence of diffusion (all *d*_*i*_ = 0), then diffusive motion of the nutrients or individuals can lead, under certain conditions, to the emergence of spatially inhomogeneous patterns. This may seem counter-intuitive at first, as diffusion is generally considered as a process that reduces spatial inhomogeneities, instead of creating them.

Stability of the homogeneous steady state in the absence of diffusion implies that the eigenvalues of the Jacobian

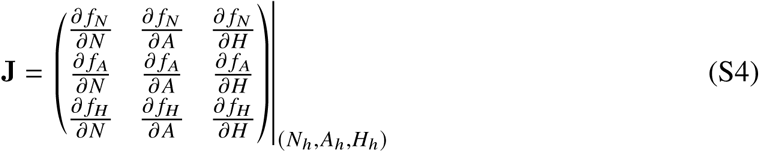

all have negative real parts. With diffusion, this steady state is destabilized by a spatially heterogeneous perturbation when the dominant eigenvalue *λ* of

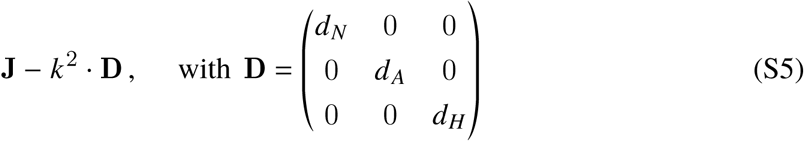

has a positive real part for some value of *k* ≠ 0. Here, *k* is the wave number of a Fourier component of the spatial perturbation. Excluding *k* = 0 simply means ignoring any spatially homogeneous component of the perturbation, as this would not contribute to the onset of pattern formation anyway. Expression (S5) yields *λ*(*k*) as functions of the wave number *k* and the task is to find diffusion constants for which *Re*(*λ*) *>* 0 for some range of *k*-values. One important observation one makes during this endeavour is that for *d*_*N*_ = *d*_*A*_ = *d*_*H*_ this never happens, i.e., a Turing instability requires the diffusion constants of the nutrients, plants, and herbivores to be different. This important result also holds for spatially discrete systems.

A simple example of a spatially discrete system (a ring of cells) was also considered by [Turing, 1952], but the general extension of his ideas to arbitrary networks of interconnected patches (or reaction sites) was provided much later [Othmer and Scriven, 1971], with some important insights being discovered only recently [Nakao and Mikhailov, 2010, Brechtel et al., 2018]. Most importantly, despite the conceptual difference between continuous space (essentially a featureless plain) and a possibly very complex network of discrete patches, the equations to be evaluated are very similar. For a network with *n* patches, instead of expression (S5), one now has to evaluate the eigenvalues of the *n* matrices

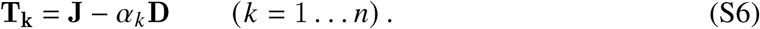

The matrices **J** and **D** are the same as before, and the wave number *k* is replaced by *α*_*k*_, the eigenvalues of the Laplacian matrix **L**. This *n* × *n*-matrix describes the structure of the network of patches and is in the simple case of undirected links of identical (unit) coupling strength constructed as follows: If two patches *i* and *j* are connected by a link (dispersal route), the elements *L*_*i j*_ and *L* _*ji*_ of **L** are set to -1, otherwise they are set to 0. Last, the diagonal elements of **L** are set to minus the sum of the off-diagonal elements of the corresponding row. For a ring of *n* patches, where each patch is linked only to its direct neighbours to the left and to the right, this leads to

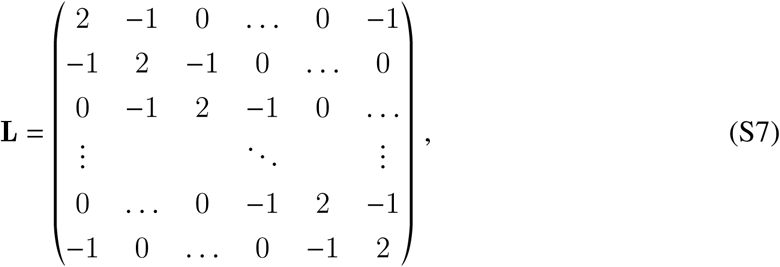

which is essentially the negative of the discretization of the Laplace operator 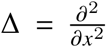 of the diffusion terms in Eqs. (S1)–(S3). The eigenvalues *α*_*k*_ of this matrix can be calculated analytically as

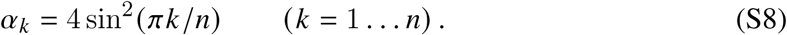

Similar to spatially continuous systems, the condition for a Turing instability is that the real part of the dominant eigenvalue of any of the *n* matrices **T**_**k**_, Eq. (S6), is positive. One therefore has to evaluate the eigenvalue spectra of at most *n* − 1 matrices to establish whether or not pattern formation can be expected for a given set of diffusion constants. One eigenvalue of the Laplacian is always 0, and symmetries in the arrangement of the patches (and thus in the Laplacian) can lead to a significant reduction in the number of distinct eigenvalues. For example, the Laplacian of a ring of patches has only *n*/2 numerically distinct eigenvalues ≠ 0.

When the attractor of the spatially homogeneous system is not a steady state (fixed point) but a limit cycle, one has to evaluate the Jacobian **J**, Eq. (S4), along this limit cycle. This makes **J** and consequently **T**_**k**_ time-dependent periodic matrices. The condition for a Turing instability is now that the absolute value of the dominant Floquet multiplier *µ* of any of the **T**_**k**_ is greater than 1. Calculating Floquet multipliers of periodic matrices is numerically more laborious than calculating eigenvalues of constant matrices, but given the small size of the matrices involved here not a major obstacle. A gentle introduction to the application of Floquet theory to ecological problems is provided by Klausmeier [2008].

### S2: Boundaries of the Turing instability

In order to establish boundaries of the diffusion constants at which a Turing instability occurs, one has to evaluate the eigenvalues of the matrices **T**_**k**_, Eq. (S6). These are given by the roots of the characteristic polynomial

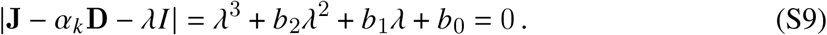

Instead of directly solving this equation for *λ*, it is sufficient to evaluate the Routh-Hurwitz stability criterion. According to this criterion, the system is unstable under diffusion if one of the following conditions is violated:

i. *b*_0_ *>* 0
ii. *b*_2_ *>* 0
iii. *b*_1_*b*_2_ − *b*_0_ *>* 0.

Condition (ii) is always met, as *b*_2_ = −*Tr* (**J**) + *α*_*k*_ (*d*_*N*_ + *d*_*A*_ + *d*_*H*_) *>* 0. This holds because *α*_*k*_ and the diffusion constants are positive and the fixed point of the non-spatial system is assumed to be stable, which implies that the trace of **J** is negative. The other two conditions can be violated for certain combinations of the diffusion constants.

To find the boundaries of the corresponding ranges of diffusion constants, we set *d*_*A*_ to a constant value and numerically solved *b*_*0*_ = 0 and *b*_1_*b*_2_ − *b*_0_ = 0 for *d*_*H*_ as a function of *d*_*N*_. Different Laplacian eigenvalues *α*_*k*_ lead to different solutions, and the eventual boundaries of the regions in the (*d*_*N*_, *d*_*H*_) plane where a Turing instability occurs are constructed as the envelope of these individual solutions. In Fig. S1 the boundaries are shown for *d*_*A*_ = 0, but they do not change much if a small, positive value for *d*_*A*_ is chosen.

**Figure S1:**
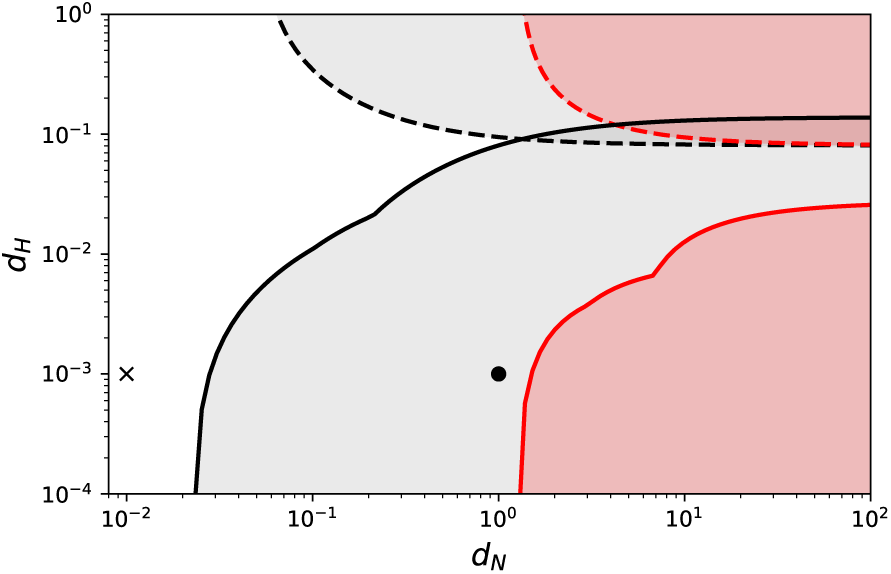
Boundaries of the Turing instability in the (*d*_*N*_, *d*_*H*_) -plane. Dashed lines denote the boundaries calculated from condition (i) of the Routh-Hurwitz stability criterion, solid lines denote the boundaries calculated from condition (iii) of the Routh-Hurwitz stability criterion. Black lines were calculated for the undefended fixed point, red lines for the defended fixed point. Self-organized pattern formation occurs in the grey- and red-shaded regions, respectively. The markers denote the combinations of *d*_*N*_ - and *d*_*H*_ -values used for Figs. 2 and 3 in the main text (x for cases 1 and 2, filled circle for cases 3 and 4).

### S3: Local Dynamics

In Fig. S2, representative time series of the dynamics of nutrient concentration *N*, biomass densities of autotrophs *A* and heterotroph *H*, the mean trait 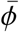, and the trait variance *v* on a single patch are shown for the defended (Fig. S2a) and undefended attractor (Figs. S2b and c). Depending on the initial conditions, the trait variance *v* can initially increase. This happens if the mean trait 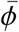 is far from its optimal value, such that the curvature of the fitness landscape 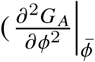,first term in the brackets in Eq. 5) is positive. Eventually, however, 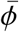 approaches its optimal value, where the curvature of the fitness landscape is necessarily negative, and *v* tends towards zero.

Fig. S3 summarises how the attractors change as a function of the maximal attack rate *a*_*max*_. On the defended attractor (Fig. S3a), the nutrient concentration, biomass densities, and mean trait value do not change much with *a*_*max*_. For values of *a*_*max*_ ≲ 1, this attractor does not exist. The undefended attractor (Fig. S3b) exists for the whole range of *a*_*max*_-values and undergoes a Hopf bifurcation at *a*_*max*_ ≈ 1.35.

**Figure S2:**
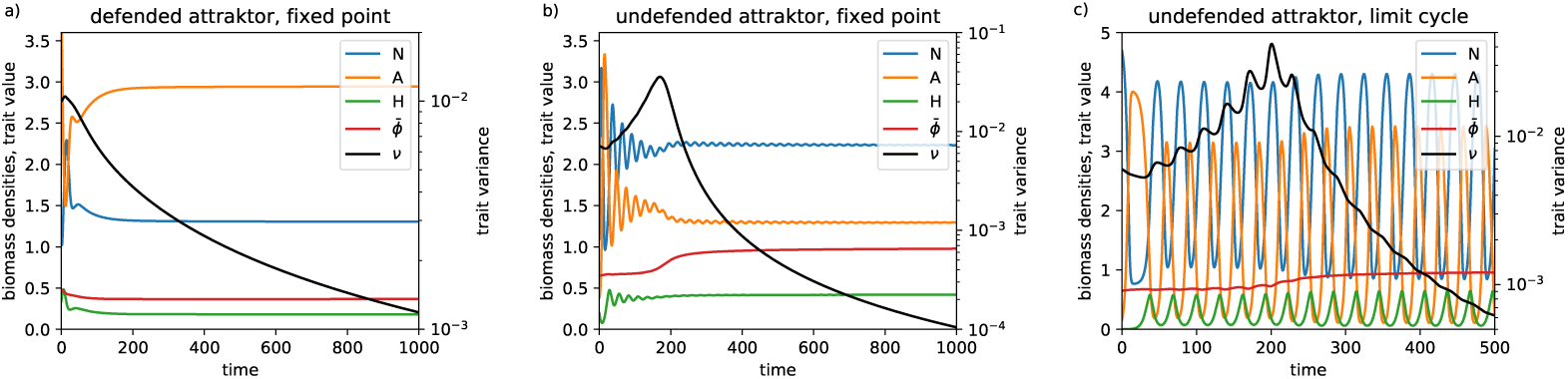
Time series of the local dynamics (single patch). a) Defended attractor as fixed point at *a*_*max*_ = 1.3. b) Undefended attractor as fixed point at *a*_*max*_ = 1.3. c) Undefended attractor as limit cycle at *a*_*max*_ = 2.0. Note that trait variance is shown on a logarithmic scale (right y-axis).

**Figure S3:**
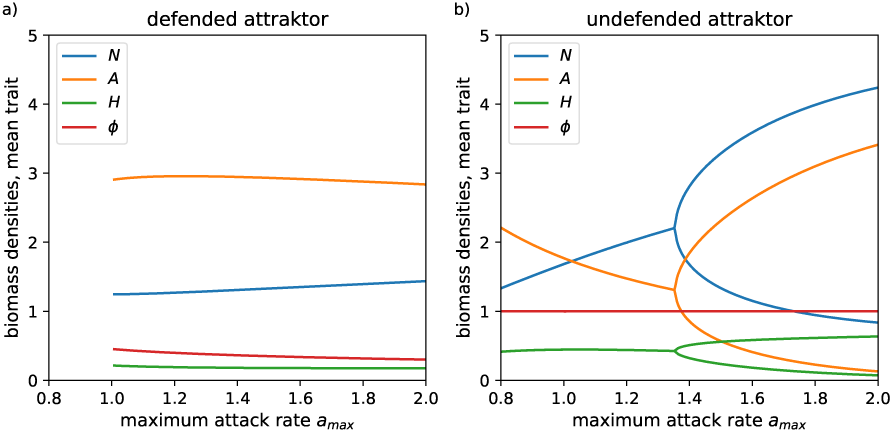
Bifurcation diagram of the local dynamics with *a*_*max*_ as bifurcation parameter. (a) defended attractor, (b) undefended attractor. The trait variance *v* is not shown because it is always at zero on the attractors.

### S4: Loss of local diversity in systems with pattern formation

When in a metacommunity all local communities are on the same attractor, local functional diversity declines towards zero even if self-organized pattern formation occurs. In Fig. S4 this is exemplarily shown for three different cases: i) all local communities on the defended fixed point (first column), ii) all local communities on the undefended fixed point (second column), and iii) all local communities on the undefended limit cycle (third column). It can be seen that in all cases different spatio-temporal patterns emerge in the biomass densities of the autotrophs (first row), but trait variance *v* always declines towards 0 (third row). In case i) the trait variance declines slowest, therefore one can still detect some trait dynamics (second row), albeit with a very small amplitude. In the other two cases, the trait variance is already at such a low level that the trait *φ* appears constant.

**Figure S4:**
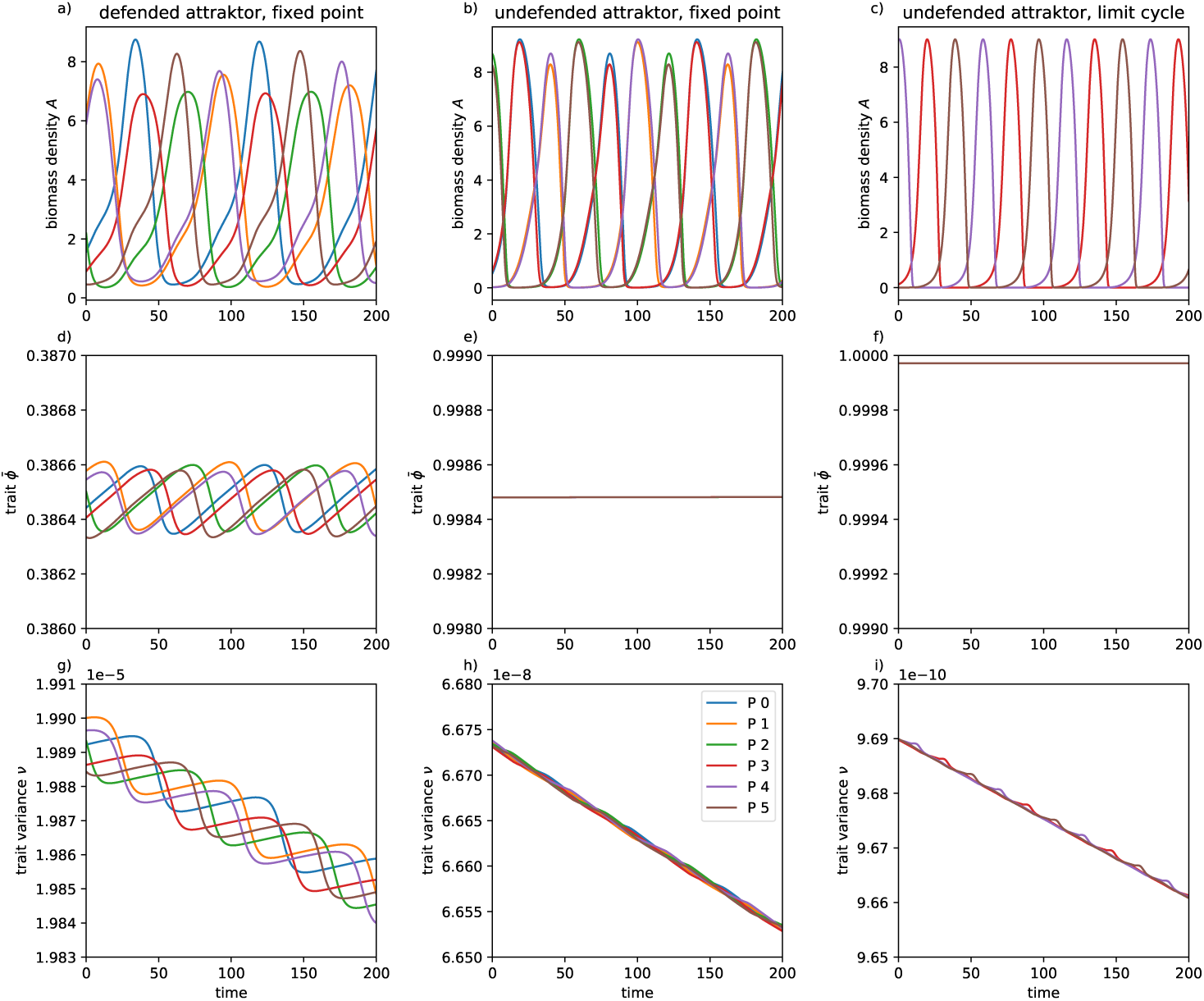
Time series of autotroph biomasses _*i*_ (top row), mean traits 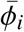 (middle row) and trait variances *v*_*i*_ (bottom row) in systems with 6 patches. Left column: all local communities on the defended fixed point (*a*_*max*_ = 1.3, *d*_*N*_ = 10), middle column: all local communities on the undefended fixed point (*a*_*max*_ = 1.3, *d*_*N*_ = 1), right column: all local communities on the undefended limit cycle (*a*_*max*_ = 2, *d*_*N*_ = 1). In all simulations *d*_*A*_ = 10^−5^, *d*_*H*_ = 10^−3^ and standard values for the other parameters.

### S5: Correlation between the distance to the Turing instability and local diversity

In this section we provide additional information regarding the correlation between the real part of the maximum dominant eigenvalue *λ* (Fig. S5a) or absolute value of the dominant Floquet multiplier *µ* (Fig. S5b), respectively, of the matrices **T**_**k**_, and the amount of local functional diversity as measured by the mean trait variance *v*. Dominant eigenvalues of **T**_**k**_ are evaluated when the undefended attractor is a fixed point, dominant Floquet multipliers are evaluated when it is a limit cycle. For these simulations, *d*_*N*_ was logarithmically varied in 20 steps between 10^−2^ and 10, *d*_*H*_ was logarithmically varied in 20 steps between 10^−4^ and 1, and *d*_*A*_ was fixed at 10^−5^. For each combination of diffusion constants 100 replicates with random initial conditions were simulated.

We evaluated the Spearman rank-correlation between Re(*λ*) or | *µ*| and the mean trait variance *v* based on the data points where the combination of diffusion constants can lead to self-organized pattern formation (i.e., for which Re(*λ*) > 0 or | *µ*| *>* 1, red points in Fig.S5). For combinations of the diffusion constants where no Turing instability occurs, calculating the correlation between Re(*λ*) or | *µ*| and the mean trait variance is obviously meaningless. For the correlation between Re(*λ*) and mean *v*, the Spearman rank-correlation correlation coefficient *r*_*S*_ = 0.86, based on 240 data points, and for the correlation between | *µ*| and mean *v*, the Spearman rank-correlation correlation coefficient *r*_*S*_ = 0.84, based on 171 data points. As is usual for simulation studies, significance is not an issue due to the high number of data points (*p <* 10^−40^ in both cases).

**Figure S5:**
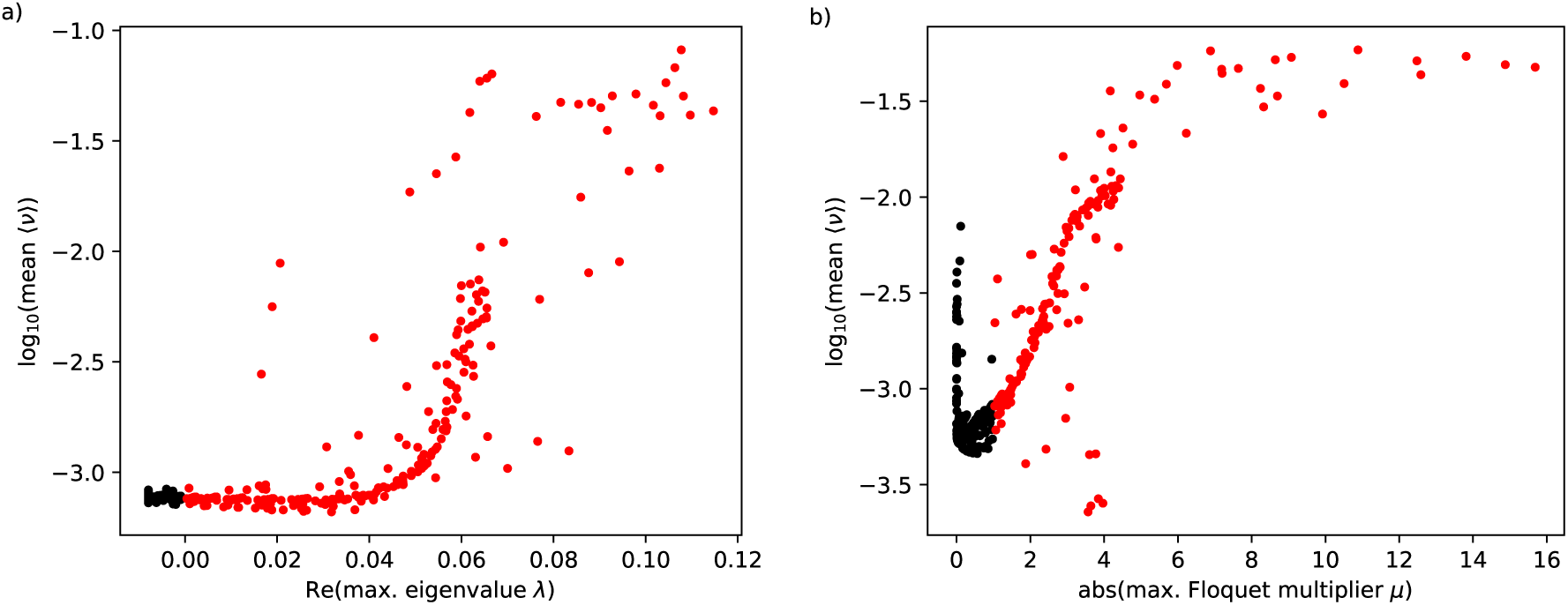
Correlation between the maximum dominant eigenvalue *λ* (left) or the maximum dominant Floquet multiplier *µ* of the matrices **T**_**k**_ and the mean trait variance *v*. For red data points, self-organized pattern formation occurs, for black data points it does not.

### S6: Sensitivity analysis

Critical parameters that might affect the amount of maintained functional diversity are those that determine the shape of the trade-off between growth rate *r* (ϕ) and attack rate *a* (ϕ), *s, b*, and *c*, as well as the number of patches, *n*.

We find that both local and regional diversity indeed depend on the shape of the trade-off (Figs. S6 - S8). However, the main result that self-organized pattern formation significantly increases local diversity and slightly decreases regional diversity is always reproduced. Except for very small patch numbers the results are almost independent of the number of patches (Fig. S9).

**Figure S6:**
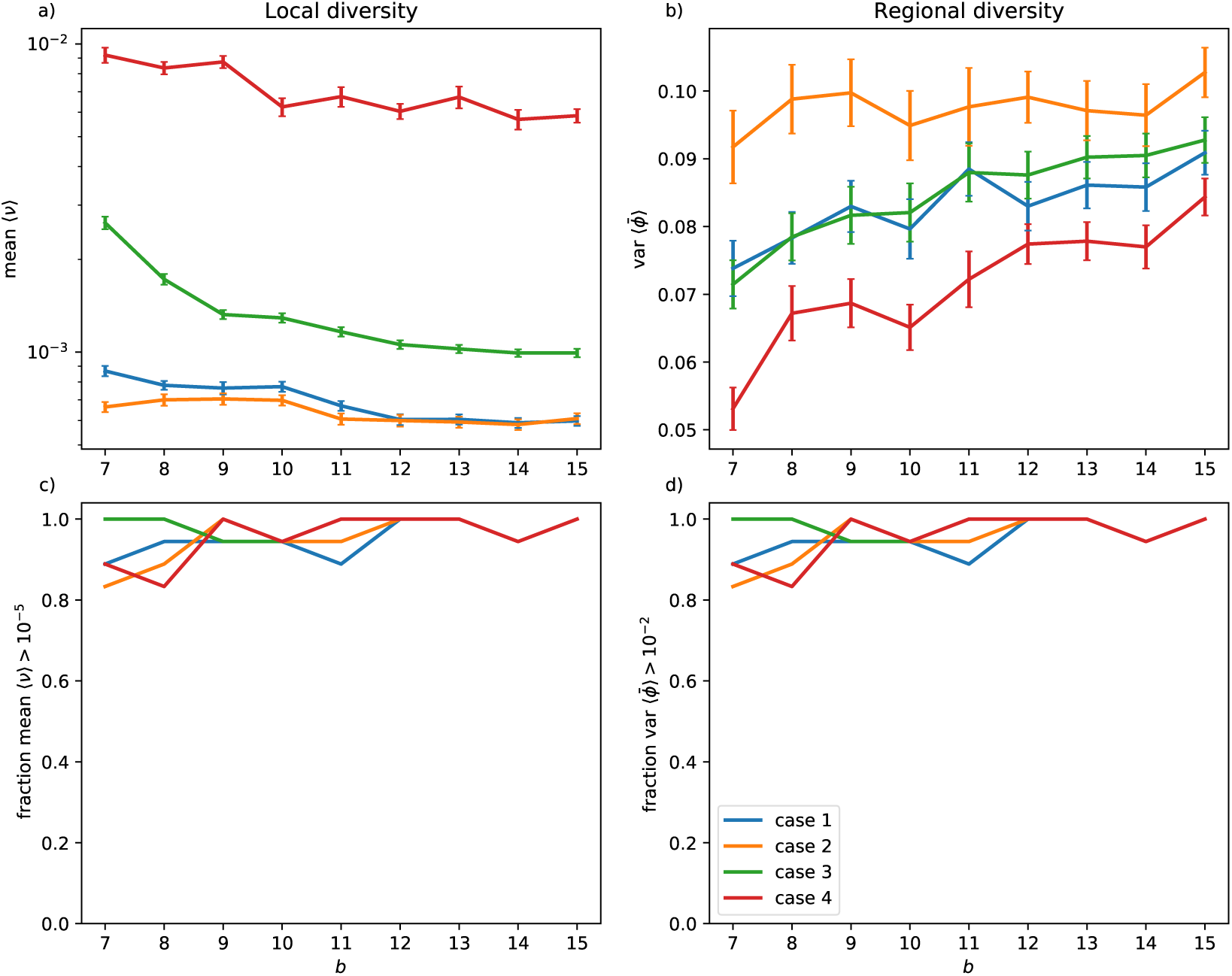
Effect of parameter *b*. In panel a) the mean trait variance *v* across 6 patches is shown as a measure of local diversity, in b) the variance of mean trait values 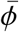 signifies regional diversity. Results were obtained as arithmetic means over 40 simulation runs with randomized initial conditions, error bars denote standard errors. In panels c) and d), the fraction of simulation runs in which local (c, mean (*v*) *>* 10^−5^) or regional 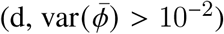 diversity were maintained, are shown.

**Figure S7:**
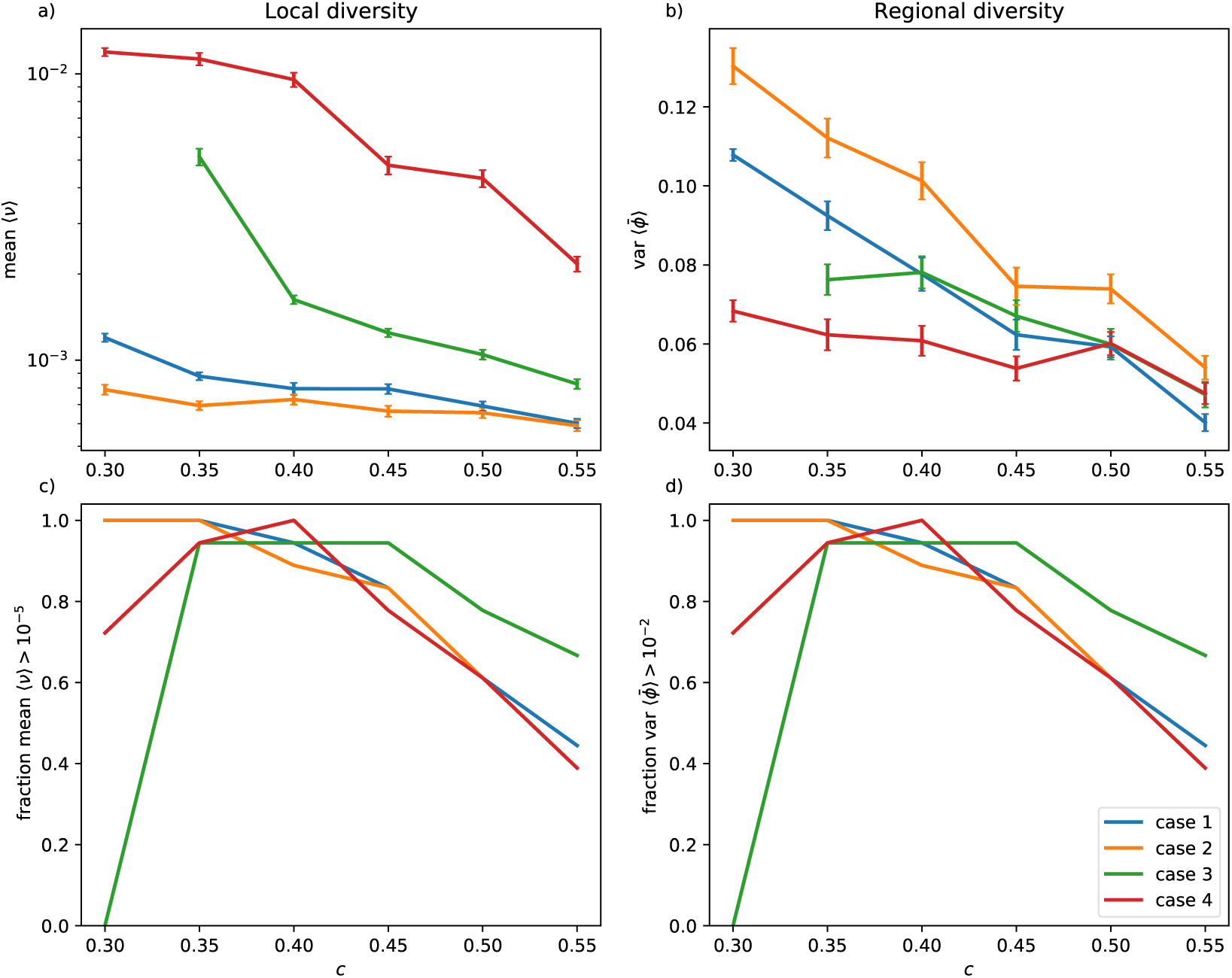
Effect of parameter *c*. In panel a) the mean trait variance *v* across 6 patches is shown as a measure of local diversity, in b) the variance of mean trait values 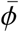 signifies regional diversity. Results were obtained as arithmetic means over 40 simulation runs with randomized initial conditions, error bars denote standard errors. In panels c) and d), the fraction of simulation runs in which local (c, mean (*v*)*>* 10^−5^) or regional 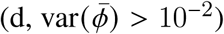 diversity were maintained, are shown.

**Figure S8:**
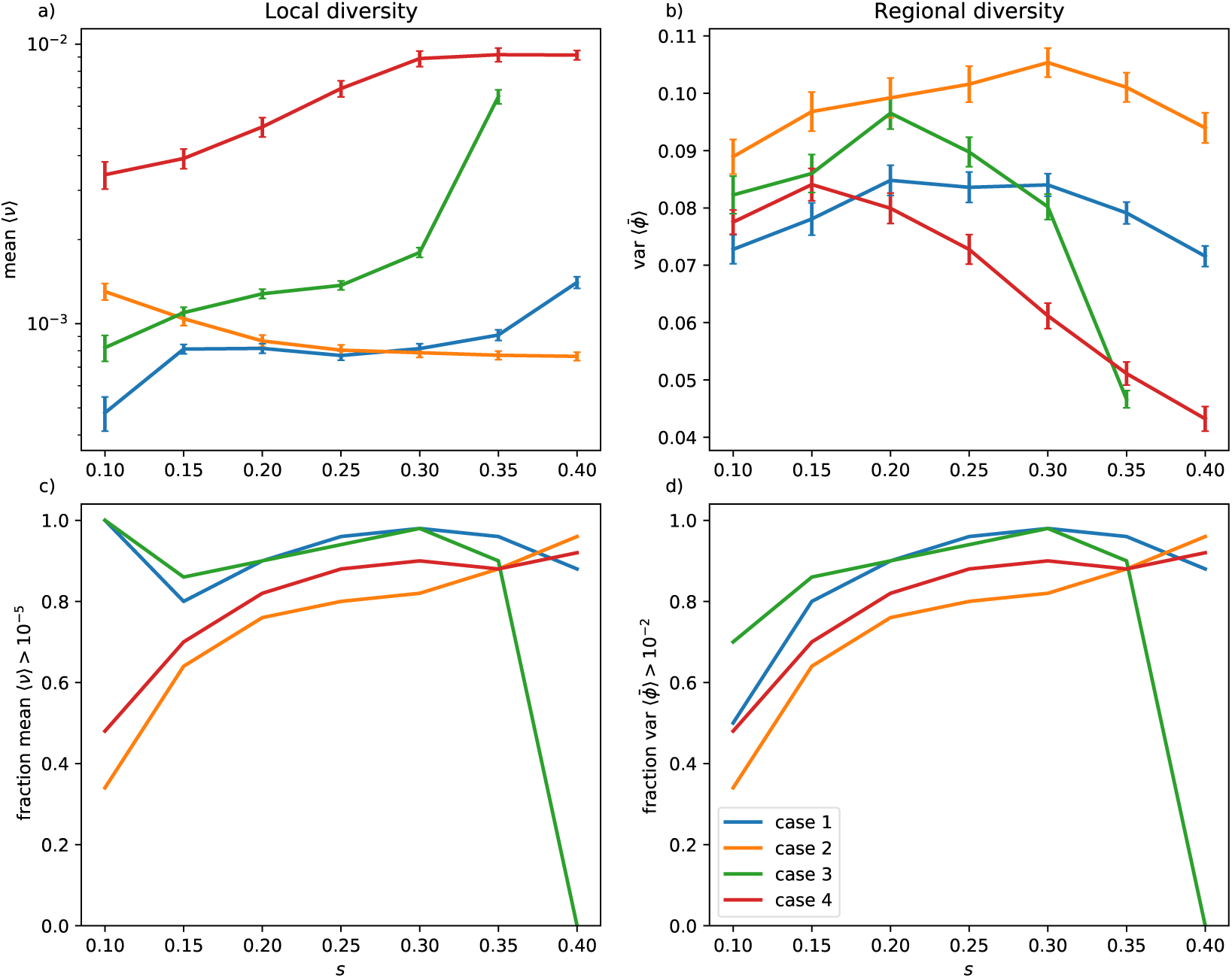
Effect of parameter *s*. In panel a) the mean trait variance *v* across 6 patches is shown as a measure of local diversity, in b) the variance of mean trait values 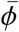 signifies regional diversity. Results were obtained as arithmetic means over 40 simulation runs with randomized initial conditions, error bars denote standard errors. In panels c) and d), the fraction of simulation runs in which local (c, mean (*v*) *>* 10^−5^) or regional 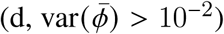 diversity were maintained, are shown.

**Figure S9:**
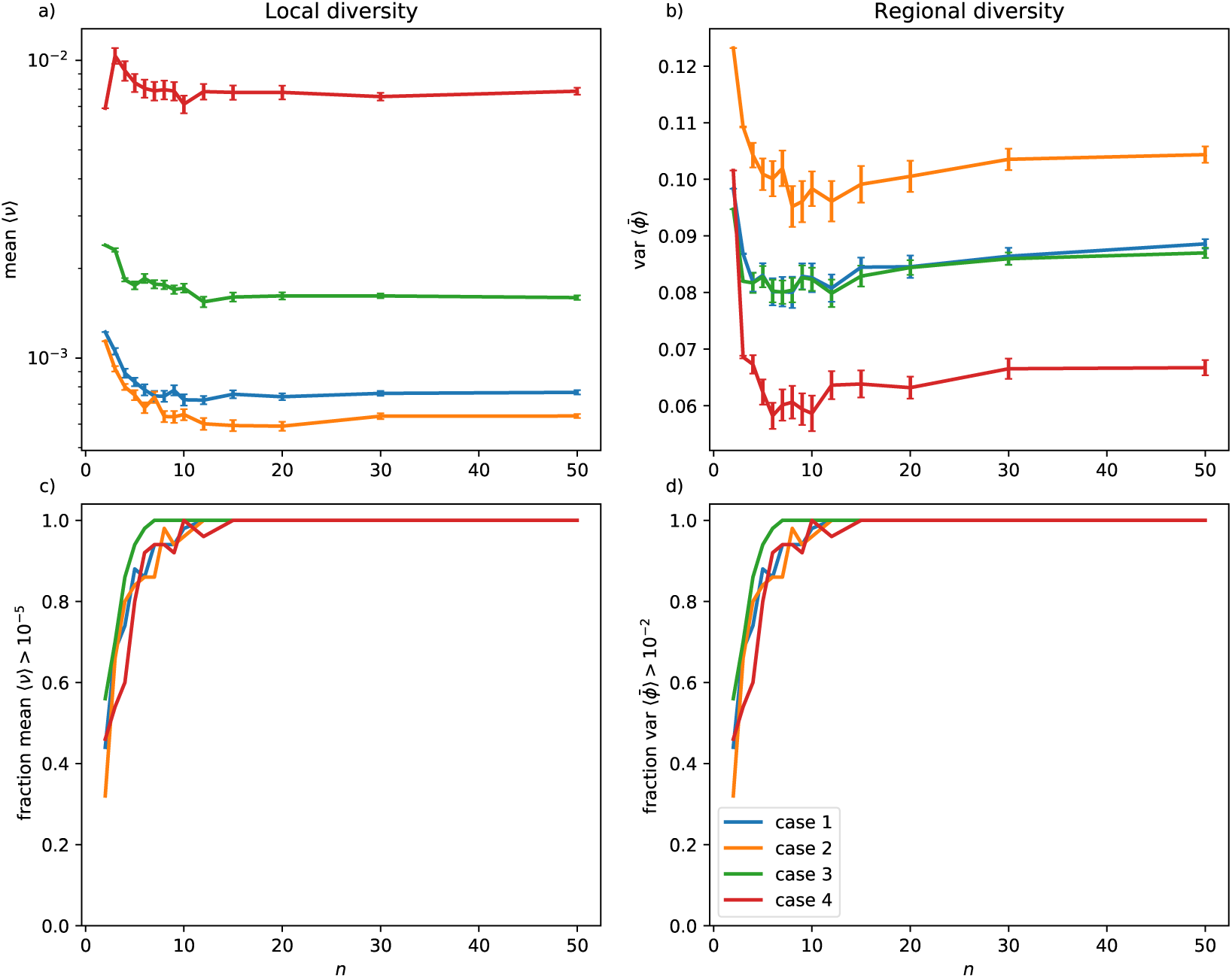
Effect of the number of patches, *n*. In panel a) the mean trait variance *v* across *n* = 2 … 50 patches is shown as a measure of local diversity, in b) the variance of mean trait values 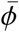 signifies regional diversity. Results were obtained as arithmetic means over 40 simulation runs with randomized initial conditions, error bars denote standard errors. In panels and d), the fraction of simulation runs in which local (c, mean(*v*) *>* 10^−5^) or regional 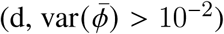a diversity were maintained, are shown.

## Notes

### Competing Interest Statement

The authors have declared no competing interest.

